# Single-molecule displacement mapping indicates unhindered intracellular diffusion of small (<~1 kDa) solutes

**DOI:** 10.1101/2023.01.26.525579

**Authors:** Alexander A. Choi, Limin Xiang, Wan Li, Ke Xu

## Abstract

While fundamentally important, the intracellular diffusion of small (<~1 kDa) solutes has been difficult to elucidate due to challenges in both labeling and measurement. Here we quantify and spatially map the translational diffusion patterns of small solutes in mammalian cells by integrating several recent advances. In particular, by executing tandem stroboscopic illumination pulses down to 400-μs separation, we extend single-molecule displacement/diffusivity mapping (SM*d*M), a super-resolution diffusion quantification tool, to small solutes with high diffusion coefficients *D* of >300 μm^2^/s. We thus show that for multiple water-soluble dyes and dye-tagged nucleotides, intracellular diffusion is dominated by vast regions of high diffusivity ~60-70% of that *in vitro*, up to ~250 μm^2^/s in the fastest cases. Meanwhile, we also visualize sub-micrometer foci of substantial slowdowns in diffusion, thus underscoring the importance of spatially resolving the local diffusion behavior. Together, these results suggest that the intracellular diffusion of small solutes is only modestly scaled down by the slightly higher viscosity of the cytosol over water, but otherwise not further hindered by macromolecular crowding. We thus lift a paradoxically low speed limit for intracellular diffusion suggested by previous experiments.

**Abstract Graphic:** 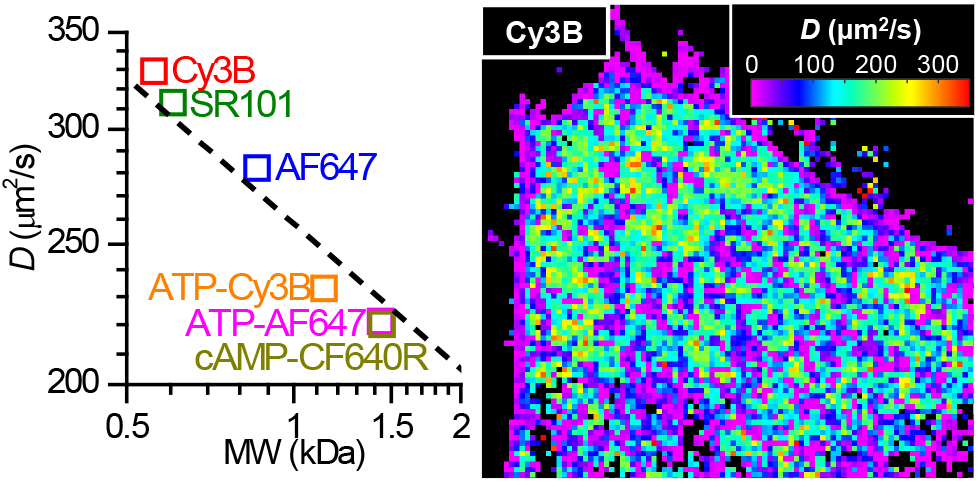

## Text

Molecular diffusion is a fundamental process of the cell that underlies key functions.^1–8^ Whereas the translational diffusion of macromolecules as proteins is routinely probed in the mammalian cell with modern live-cell tagging and diffusivity measurement approaches, the intracellular diffusion of unbound small (<~1 kDa) solutes has been more difficult to elucidate.

On the labeling side, as small fluorescence probes cannot be expressed in the cell [typical fluorescence proteins (FPs) are ~30 kDa], they often need to be delivered exogenously. While lipophilic fluorescent dyes pass the plasma membrane, their strong lipid interactions make them mainly report on the slow (diffusion coefficient *D* <~3 μm^2^/s) diffusion of intracellular membranes.^9,10^ Conversely, hydrophilic dyes are typically membrane impermeable. Capping the polar functional groups, *e.g.*, with acetoxymethyl ester or acetate ester, enables intracellular delivery with subsequent de-capping, yet dyes delivered this way may be sequestered in vesicles.^11^ More generally, independent of the loading method, a fraction of the delivered probes may be bound to different intracellular structures,^12–14^ leading to reduced apparent diffusivity.

On the measurement side, the very fast diffusion (*D* >~200 μm^2^/s; see below) of unbound small molecules is difficult to quantify: the range is orders of magnitude higher than what is workable with typical single-molecule tracking,^15,16^ while also causing large uncertainties for other common microscopy-based approaches as fluorescence recovery after photobleaching (FRAP) and fluorescence correlation spectroscopy (FCS).^1,5,7,17^ It is even more difficult to map out spatial heterogeneities in intracellular diffusion, and the global readout may be biased by the locally bound molecules. Separately, whereas incell NMR provides alternative means to probe molecular diffusion,^18^ ensemble results are collected from many cells; cell confinement effects and signal from the extracellular space, together with the limited signal-to-noise ratio, complicate data interpretation.

With the above challenges, our current model of how small solutes diffuse in the cell^2,6^ has often focused on an early FRAP study^19^ in which the acetoxymethyl ester-loaded 520 Da dye BCECF is found to diffuse four-time slower in the mammalian cytoplasm versus *in vitro* (*D*_cyto_/*D*_water_ ~0.25). While this large reduction in *D* sets an alarming speed limit for the intracellular diffusion of small molecules, it contrasts with the modest (~20-40%) drop in intracellular rotational diffusivity of the same dye, which suggests only a slight increase in the cytoplasm viscosity over water.^20^ Although this paradox may be reconciled by considering hindered translational diffusion *via* collision with cell solids,^2,19^ similar measurements on intracellularly expressed GFP show slightly less suppressed *D* (*D*_cyto_/*D*_water_ ~0.31),^21^ even as stronger macromolecular crowding-induced hindrances may be expected for the much larger GFP. Later experiments have reported scattered intracellular diffusivities for different small solutes,^22–27^ and with the above-noted limitations in cell loading and measurement, have not resolved the *D*_cyto_/*D*_water_ ~0.25 speed-limit paradox.

Here we overcome previous challenges to quantify and map out the intracellular diffusivity of small solutes by integrating several recent advances. For intracellular delivery, we exploit the discovery that astrocytes actively uptake the hydrophilic dye sulforhodamine 101 (SR101),^28,29^ and then move to other small solutes *via* our recently devised graphene-based *in situ* electroporation.^30^ For *D* measurement, we harness our recent development of single-molecule displacement/diffusivity mapping (SM*d*M),^31^ which has previously enabled the super-resolution *D* mapping of ~30 kDa FPs in live cells at *D*~20 μm^2^/s. Extending SM*d*M to unbound small molecules, we here quantify *D* values of >300 μm^2^/s *in vitro* and >200 μm^2^/s in mammalian cells. We thus show that for multiple water-soluble dyes and dye-tagged nucleotides, the intracellular diffusion is dominated by large regions of unhindered diffusion with *D* ~60-70% of the diffusion *in vitro*.

In SM*d*M, wide-field images are recorded for diffusing single molecules with a typical EM-CCD camera. Stroboscopic illumination is employed to mitigate single-molecule motion-blur,^32^ and the stroboscopic pulses are applied in tandem across paired camera frames, so that single-molecule displacements are recorded for time separations substantially shorter than the camera frame time (**Figure 1a** and **Figure S1**).^31^ The paired excitation pulses across tandem frames further leave ample time between the anti-paired pulses for molecules outside the illuminated area to diffuse in, thus creating a mechanism to dynamically exchange and replenish molecules during imaging and permitting the use of conventional, non-photoswitchable fluorophores in SM*d*M.^31,33^

**Figure 1.**
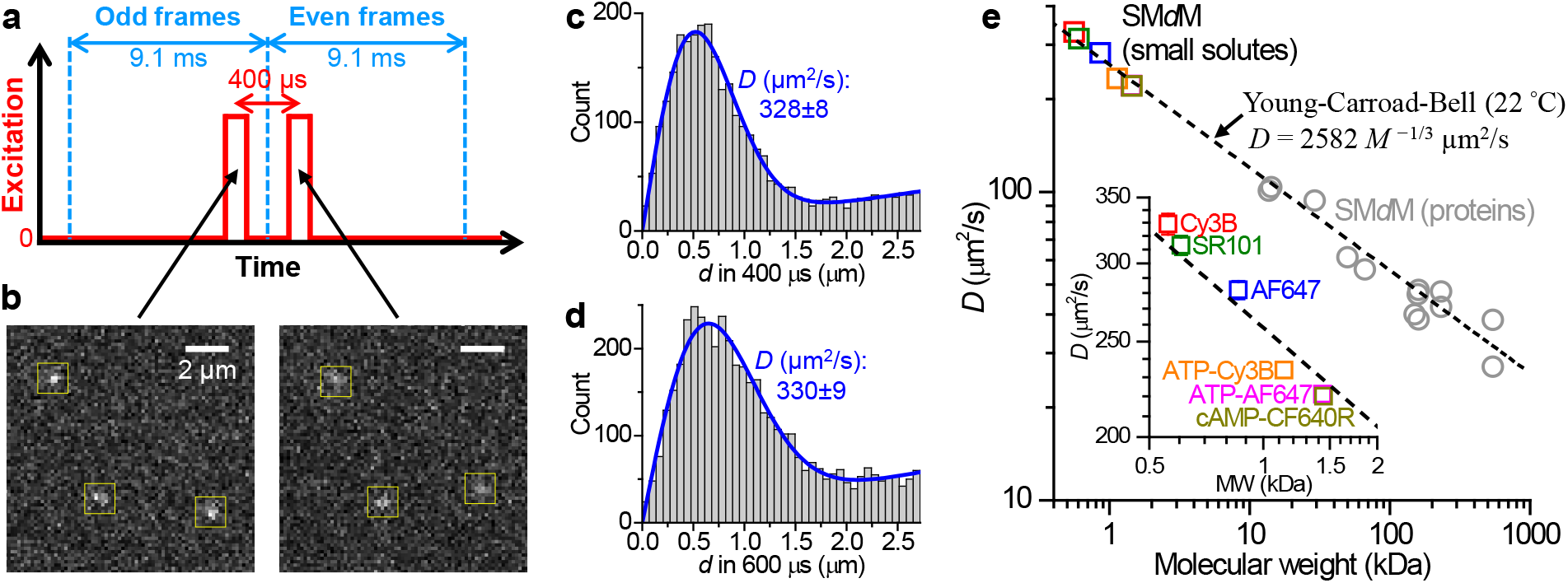
SM*d*M quantifies the fast diffusion of small solutes *in vitro*. (a) Schematics: To detect the fast motion of small solutes, tandem stroboscopic excitation pulses of 200 μs duration are applied across paired camera frames at a Δ*t* = 400 μs center-to-center separation. (b) Example single-molecule images captured in a tandem frame pair, for Cy3B freely diffusing in the PBS buffer at room temperature. (c,d) Typical distributions of the recorded singlemolecule displacements for two experiments carried out at Δ*t* = 400 μs (c) and 600 μs (d), respectively, for Cy3B in PBS. Blue lines: maximum likelihood estimation (MLE) fits to our model, yielding diffusion coefficients *D* of 328±8 and 330±9 μm^2^/s, respectively. (e) Summary of the SM*d*M-measured *D* values of 6 different small solutes in PBS (color squares), as well as 13 protein samples we previously reported (gray circles),^34^ plotted as a function of molecular weight on a logarithmic scale. Dash line: the Young-Carroad-Bell model at 22 °C: *D* = 2582 *M*^-1/3^ μm^2^/s, *M* being the molecular weight in Da. Inset: zoom-in of the small-molecule data. Error bars are shown for MLE uncertainties when larger than the symbol size.

We previously worked with a typical center-to-center pulse separation of Δ*t* = 1 ms when probing the intracellular diffusion of FPs.^31^ Here for the fast-diffusing small solutes, we dropped *Δt* down to 400 μs with the duration of each pulse *τ* accordingly reduced to 200 μs (**Figure S1**), and we focused on bright fluorescent dyes.^35,36^ Good single-molecule images were obtained, as shown in **Figure 1b** for the free diffusion of the smallest (and thus fastest) probe examined in this work, Cy3B (560 Da), in the phosphate-buffered saline (PBS). Accumulating the resultant 400-μs displacements calculated from the super-localized positions from many paired frames yielded distributions (**Figure 1c**) well fitted by our model based on normal diffusion plus a background term for mismatched molecules (Methods).^31^ Relaxing Δ*t* to 600 μs led to a broader distribution of single-molecule displacements and a slight increase in the background (**Figure 1d**). Nonetheless, similar *D* values of ~330 μm^2^/s were obtained for both experiments, comparable to that previously measured for fluorescent probes of similar sizes.^37,38^ Note that as PBS and other common buffers have viscosities near-identical to water, we collectively discuss diffusivity in them as typical “*in vitro*” values. To assess whether SM*d*M could correctly quantify impeded diffusion, we further examined Cy3B diffusion in PBS containing varying amounts of glycerol. We thus found that as glycerol addition increased the solution viscosity, the SM*d*M-measured diffusivity decreased accordingly following the expected trend over a wide range (**Figure S2**), down to ~20% of its initial value at 50% glycerol.

**Figure 1e** and inset summarize the SM*d*M-determined *D* in PBS for the 6 different small solutes we examined in this work. A notable drop in *D* was observed for the larger molecules. Moreover, as we plotted the *D* values against the molecular weight (*M*) on a logarithmic scale and further added in our previous *in vitro* SM*d*M results of 13 protein samples,^34^ they all agreed reasonably well with the Young-Carroad-Bell Model^39^ without any fitting or scaling (**Figure 1e**). This agreement was not expected *a priori* (as the Model was summarized from >~3 kDa proteins), yet reasonable in hindsight: few-heteroatom organic compounds may have densities comparable to that of protein. Their molar volumes, and hence *D*, may thus follow similar trends versus *M* as proteins.

After validating SM*d*M’s applicability to the fast diffusion of small solutes, we started cell experiments with the special case of the 607 Da, yellow-excited dye SR101 (**Figure 2b**). While being water-soluble and generally cell-impermeable, SR101 is actively uptaken by astrocytes.^28,29^ Thus, by incubating cultured primary rat hippocampal astrocytes with an SR101-containing medium, the dye was readily loaded into the cells. SM*d*M registered good single-molecule images (**Figure 2a**). As the camera recorded at ~110 frames per second, time separations between the anti-paired pulses were >35 times longer than between the paired pulses (**Figure S1**). Consequently, whereas single-molecule displacements were tracked between the paired frames (*e.g.*, Frames 1-2 and Frames 3-4 in **Figure 2a**), molecules in the view were well exchanged via diffusion between the anti-paired pulses (*e.g.*, Frames 2-3 in **Figure 2a**). As we illuminated and imaged an ~1 μm thin layer at the bottom of the cell (Methods), diffusional exchanges enabled sustained single-molecule imaging with only a modest drop in the molecular count over >7×10^4^ frames (**Figure 2c**). Spatially binning the accumulated single-molecule displacements onto 320 nm-sized grids enabled local fitting^31,33^ to extract *D* values and construct *D* maps (**Figure 2d**).

**Figure 2.**
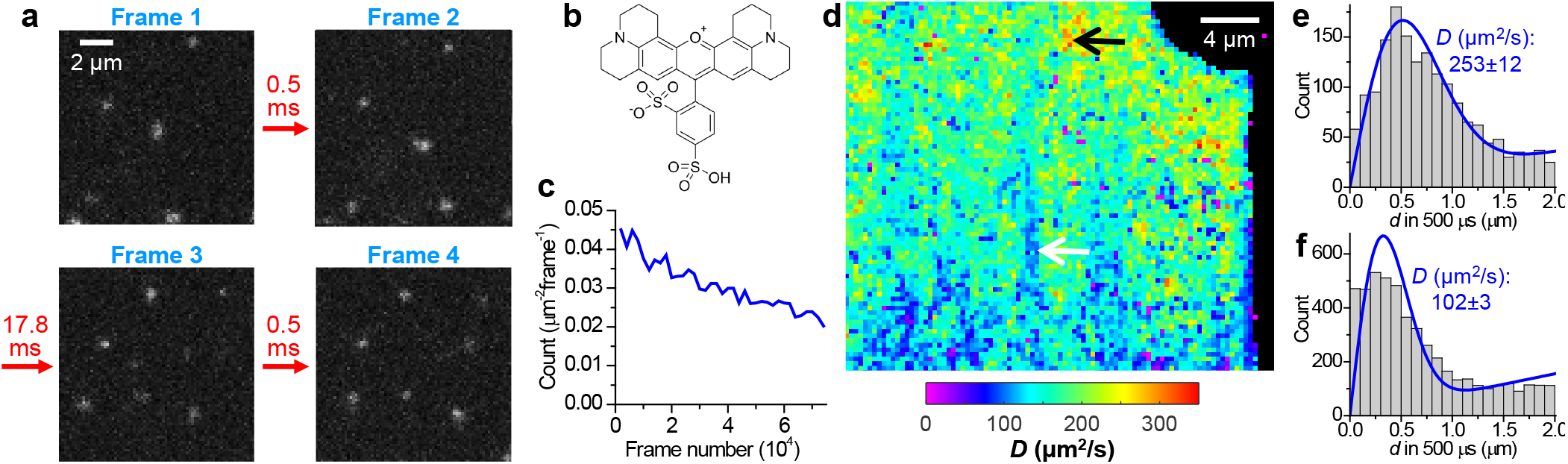
SM*d*M of the water-soluble dye SR101 in primary rat hippocampal astrocytes. (a) Four consecutive frames of the SM*d*M-recorded single-molecule images of SR101 diffusing inside an astrocyte, with excitation pulse length *⊤* = 200 μs and center-to-center separation *Δt* = 500 μs. (b) Chemical structure of SR101. (c) Count of detected single molecules per μm^2^ per frame over the recording of 7.5×10^4^ frames. (d) Color-coded SM*d*M *D* map of SR101 in the cell. (e,f) Local distributions of the recorded 500-μs single-molecule displacements for ~1 μm^2^ regions of fast (e) and slow (f) diffusion in the cell, as pointed to by the black and white arrows in (d), respectively. Blue lines: MLE fits to our model, yielding *D* of 253±12 and 102±3 μm^2^/s, respectively.

We thus found that for vast regions of the astrocyte cytoplasm, SR101 diffused at *D*>~200 μm^2^/s (**Figure 2d**), >~60% of that in PBS (**Figure 1e**). As a control, we compared SM*d*M under the same excitation laser for astrocytes not loaded with SR101 but expressing the mEos3.2 FP, and observed *D* ~20 μm^2^/s (**Figure S3**), comparable to our previous results in mammalian cells.^31^ For the intracellular diffusion of SR101, the fastest regions reached *D*~250 μm^2^/s (**Figure 2de**), ~80% of that in PBS. Meanwhile, micrometer-sized structures of lower *D* were also noted (**Figure 2d**), where the apparent *D* dropped to ~100 μm^2^/s and the distribution was not well fit by our model based on single-component Gaussian diffusion (**Figure 2f**), indicative of population heterogeneity, *e.g.*, a fraction interacting with intracellular organellar membranes. As such interactions may be dynamic and complex, we did not attempt to fit the single-molecule displacements with multiple components; the overall smaller displacements were captured by the imperfect single-component fit as low nominal *D* values.

Although SR101 is easily loaded into astrocytes, this strategy cannot be generalized to other molecular species or cell types. To deliver additional small solutes into mammalian cells, we harnessed our recent development of graphene-based *in situ* electroporation.^30^ By coating the glass coverslip with graphene, a conductive monolayer of carbon atoms, the application of voltage pulses across the graphene-coated substrate and a counter electrode (**Figure 3b**) enables electroporation and fluorescent-probe delivery for adherent cells while maintaining full compatibility with single-molecule imaging.

**Figure 3.**
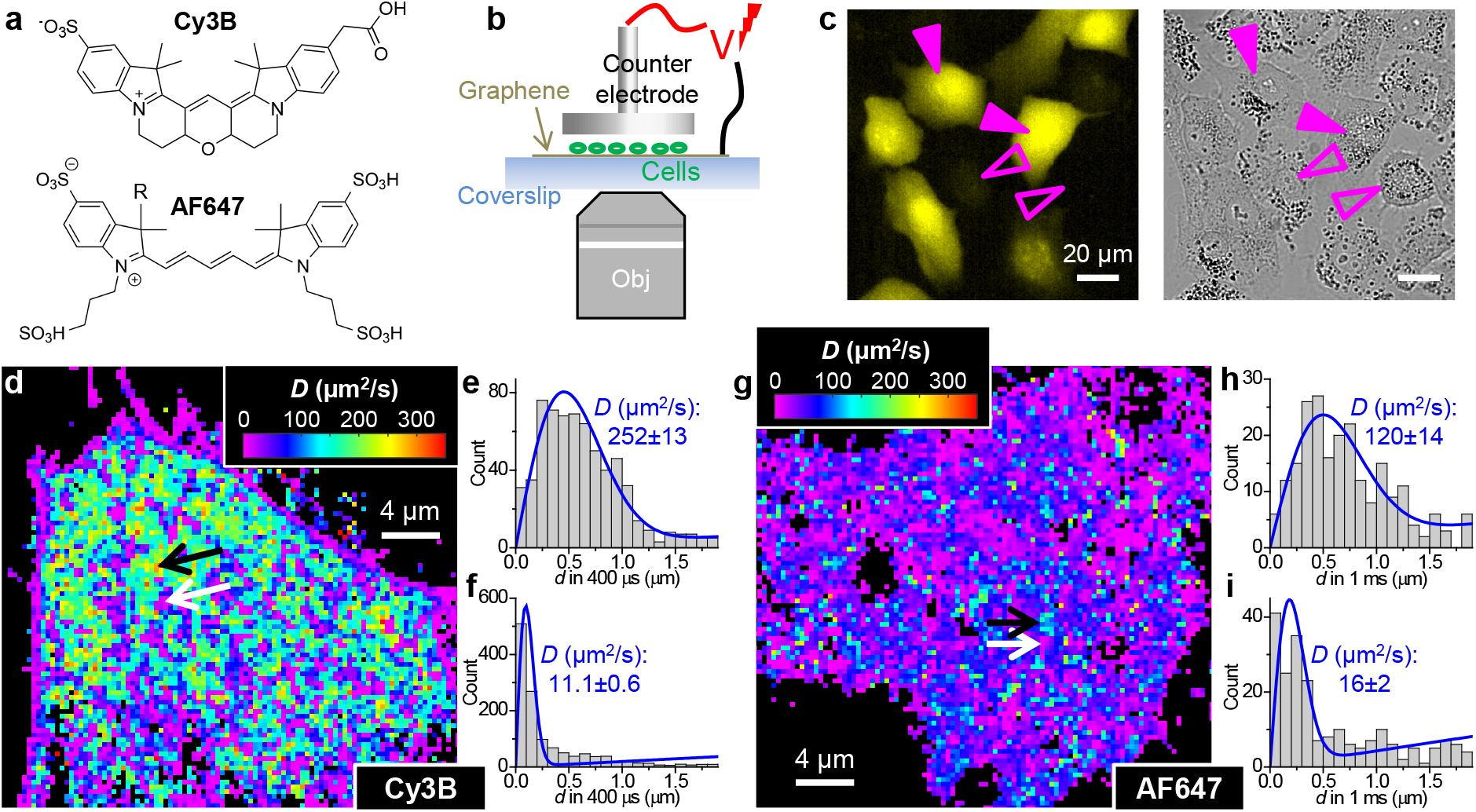
SM*d*M of Cy3B and AF647 dyes delivered into the A549 epithelial cell line via graphene-based electroporation. (a) Molecular structures of the two dyes. (b) Schematics: electroporation by applying a voltage pulse between the graphene-coated coverslip substrate and a counter electrode. (c) Example fluorescence image of Cy3B after electroporation loading into A549 cells (left), vs. bright-field image of the same sample (right). Filled and hollow arrowheads point to loaded and unloaded cells in the same sample, respectively. (d) SM*d*M *D* map of the delivered Cy3B in an A549 cell. (e,f) Local distributions of single-molecule displacements for ~1 μm^2^ regions of fast (e) and slow (f) diffusion in the cell, as pointed to by the black and white arrows in (d), respectively. (g) SM*d*M *D* map of the delivered AF647 in an A549 cell. (h,i) Local distributions of single-molecule displacements for ~1 μm^2^ regions of relatively high (h) and low (i) diffusivities. Blue lines in (e,f,h,i): MLE fits to our model, with resultant *D* values and uncertainties marked in each plot.

We start with two water-soluble fluorescent dyes, the green/yellow-excited Cy3B and the red-excited Alexa Fluor 647 (AF647) (**Figure 3a**). Both dyes are bright and widely used in single-molecule microscopy,^35,36^ and exhibit no appreciable interactions with lipid bilayers^12^ so may be less trapped by intracellular membranes. Graphene-based electroporation successfully delivered the dyes into A549 human epithelial cells (**Figure 3c**).

SM*d*M unveiled contrasting intracellular diffusion behaviors for the two dyes. The 560 Da Cy3B exhibited vast regions of *D*>~200 μm^2^/s (**Figure 3d**), >~60% of that in the PBS, and the faster regions reached *D*~250 μm^2^/s (**Figure 3de**), ~75% of that in the PBS (**Figure 1c-e**). Together with our above SR101 results in astrocytes (**Figure 2**), these observations indicate that small solutes may diffuse unhindered in the cytoplasm, only being mildly slowed by the ~20-40% higher viscosity of the cytosol over water. Meanwhile, we also observed (sub)micrometer-sized features with low nominal *D* of ~10 μm^2^/s and displacement distributions not well fitted by our single-component model (**Figure 3df**), indicative of substantial binding to intracellular structures, *e.g.*, organellar membranes. These results underscore the importance of spatially resolving the local diffusion behavior, so that unhindered diffusion can be separated from the locally bound molecules, as opposed to a mixed global readout that may bias the result to an artificially low *D*. In contrast, the 959 Da AF647 showed generally slow diffusion in the cell, with vast regions exhibiting nominal *D* <~50 μm^2^/s (**Figure 3gi**), <~18% of that in the PBS (**Figure 1e**). Micrometer-sized domains of faster diffusion were also observed, where *D* reached up to ~120 μm^2^/s (**Figure 3gh**), ~40% of that in PBS (**Figure 1e**).

To clarify whether the substantially more hindered intracellular diffusion of AF647 could be due to its larger size over Cy3B, we next examined the intracellular diffusion of Cy3B- and AF647-tagged adenosine triphosphate (ATP). SM*d*M showed that the 1136 Da ATP-Cy3B, after being loaded into A549 cells *via* graphene-based electroporation, diffused at *D*~120 μm^2^/s over vast regions (**Figure 4a**), ~50% of that in PBS (**Figure 1e**). Thus, the intracellular diffusion of ATP-Cy3B is less hindered than that of AF647 despite its larger size. In contrast, the intracellular diffusion of the 1434 Da ATP-AF647 (**Figure 4b**) was substantially slower than in the PBS, with vast regions exhibiting *D* <~30 μm^2^/s (~15% of in PBS) and sporadic domains reaching ~60 μm^2^/s (~30% of in PBS), analogous to what we observed for the AF647 dye itself (**Figure 3g**). Previous work has shown that AF647 tends to cluster up in mammalian cells,^13,14,40^ which may account for its significant slowdown in diffusion.

**Figure 4.**
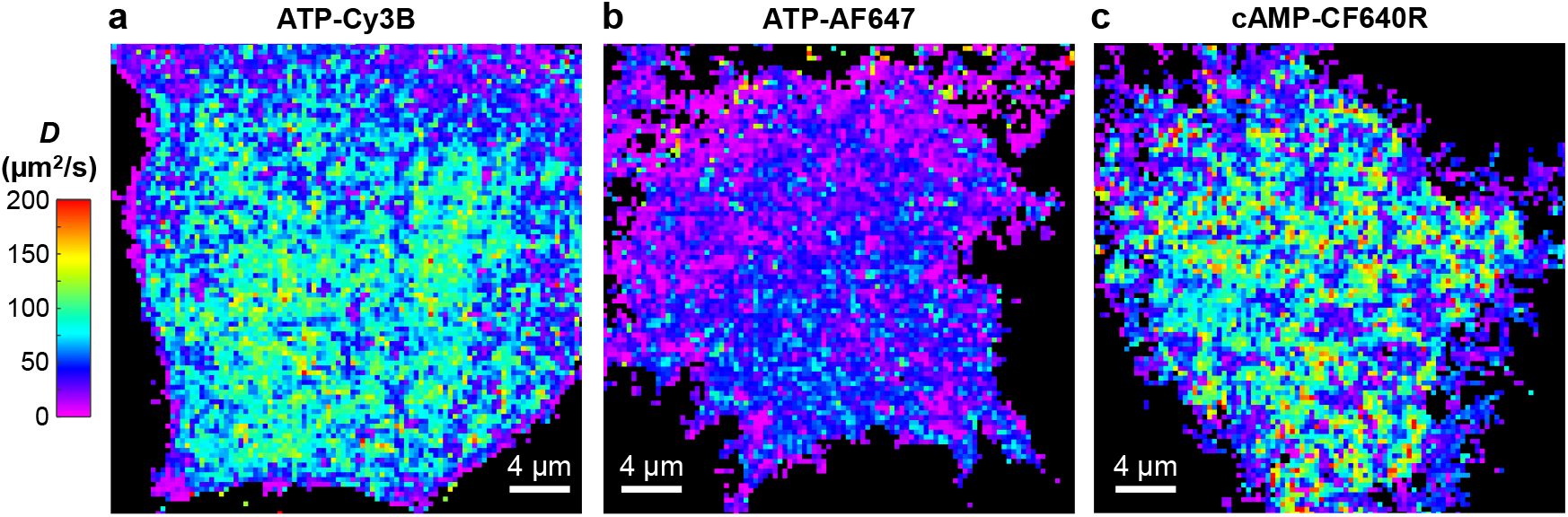
SM*d*M *D* maps of graphene-electroporation-delivered ATP-Cy3B (a), ATP-AF647 (b), and cAMP-CF640R (c) in A549 cells. Note the different *D* color scales versus Figures 2 and 3 for the slower diffusion of the larger molecules.

To further substantiate AF647 as the outlier, we examined CF640R, a different red-excited dye that has shown minimal aggregations in live cells.^13,14^ With a 1445 Da cyclic adenosine monophosphate (cAMP) conjugate, near-identical in size to ATP-AF647, we observed fast diffusion of *D*~130 μm^2^/s over vast regions in the A549 cytoplasm (**Figure 4c**), ~60% of that in PBS.

Together, by utilizing two new strategies to deliver small solutes into mammalian cells and pushing the working *D* range of SM*d*M to >300 μm^2^/s, we have unveiled the intracellular diffusion patterns for several hydrophilic dyes and their nucleotide conjugates. We thus showed that for these 560-1450 Da solutes, *D* values ~60-70% of that in the PBS could be reached in vast regions of the cell, while also noting substantial local slowdowns at the sub-micrometer scales that emphasize the importance of *D* spatial mapping. The outlying results of AF647 with substantially reduced intracellular *D* further underscore potential artifacts that may be caused by the label. As the dyes were larger in size than the tagged nucleotides examined in this work, diffusion properties were likely dominated by the former.

The *D*_cyto_/*D*_water_ ~60-70% result suggests that the intracellular diffusion of small solutes is only modestly reduced by the ~20-40% higher viscosity of the cytosol over water, but otherwise not further hindered by macromolecular crowding as seen with FPs.^1,21^ While this result contrasts with the general notion that small solutes diffuse at *D*_cyto_/*D*_water_ ~25% based on early FRAP measurements on acetoxymethyl ester-loaded BCECF,^2,6,19^ NMR has indicated unhindered diffusion of the 507 Da ATP in the rat skeletal muscle,^22^ and fluorescent reporters have implied fast diffusion of the 329 Da cAMP and 345 Da cyclic guanosine monophosphate in mammalian cells consistent with unhindered diffusion.^41^ Unhindered diffusion is plausible for small solutes: proteins and other intracellular macromolecular crowders are often substantially larger than 1 kDa, and it is likely that they do not act as effective diffusion barriers for small solutes. As the cytoplasmic transport of small solutes mainly relies on passive diffusion, lifting the previously suggested, paradoxical speed limit could carry significant implications.

## Materials and Methods

### Fluorescent probes

SR101 was from Sigma-Aldrich (S7635). Cy3B and AF647 free dyes were prepared by hydrolyzing the corresponding NHS (
*N*-hydroxysuccimidyl) esters (PA63101, Cytiva, and A200006, Invitrogen) in 0.1 M sodium bicarbonate overnight. ATP-AF647 was from Invitrogen (A22362). cAMP-CF640R was from Biotium (#00037). ATP-Cy3B was conjugated by reacting EDA-ATP (A 072, BIOLOG) with Cy3B-NHS ester at a high excess ratio (1:80). Thin-layer chromatography confirmed negligible unconjugated Cy3B in the product.

### Optical setup for SM*d*M

SM*d*M was performed on a setup^31^ based on a Nikon Ti-E inverted fluorescence microscope, but with the addition of a 642-nm laser with pulse control. Briefly, 561 nm (OBIS 561 LS, Coherent, 165 mW) and 642 nm (Stradus 642, Vortran, 110 mW) lasers were focused at the back focal plane of an oil-immersion objective lens (CFI Plan Apochromat Lamda 100×, NA = 1.45) to enter the coverslip-sample interface slightly below the critical angle, thus illuminating a few micrometers into the sample. The illuminated area was ~40 μm in diameter, yielding ~10 kW/cm^2^ peak power density. Single-molecule images were recorded in the wide field with an EM-CCD camera (iXon Ultra 897, Andor) that ran continuously at 110 frames per second. A multifunction I/O board (PCI-6733, National Instruments) detected the exposure timing signal of the camera and accordingly modulated the 561 nm (for excitation of SR101 and Cy3B) and 642 nm (for excitation of AF647 and CF640R) lasers. Paired pulses were thus applied across tandem camera frames (Figure 1a). Typical center-to-center separations between the paired pulses were Δ*t* = 400-1000 μs, with the pulse durations τ = Δ*t*/2. The timing waveforms were confirmed by a GW Instek GDS-1054B oscilloscope (**Figure S1**). The tandem excitation scheme was repeated ~10^4^ times for each SM*d*M run. As we illuminated a few micrometers into the sample, we imaged and photobleached a thin layer at the bottom. Diffusional exchange with deeper parts of the sample enabled the stochastic replenishment of single molecules in the view for sustained imaging over long recording times (**Figure 2ac**), a process facilitated by the ample time (~18 ms) between the anti-paired pulses in our excitation sequences.

### SM*d*M in PBS

25-mm diameter, #1.5 coverslips were treated in a heated 3:1 H_2_SO_4_:H_2_O_2_ (30%) mixture for 45 minutes, rinsed with Milli-Q water, and then dried using N2 gas. To reduce out-of-focus backgrounds due to fluorescent probes at the coverslip surface, the coverslips were functionalized with 10 mg/mL methoxy PEG silane (5 kDa, M-SLN-5000, JenKem Technology) in 95% ethanol/water for 30 min, and rinsed and sonicated for 5 min in Milli-Q water. Fluorescent probes were diluted to ~100 pM in PBS (14190144, Gibco), and SM*d*M was performed ~2 μm away from the glass surface in the solution.

### SM*d*M of SR101 in primary rat hippocampal astrocytes

Primary rat hippocampal astrocytes were from BrainBits and cultured on polylysine-coated #1.5 glass coverslips (GG-18-1.5-PDL, Neuvitro) in the NbAstro medium following the recommended protocols. For SR101 loading, the sample was incubated in the NbAstro medium with 100 nM SR101 for 20 min, followed by rinsing with PBS for three times. Previous studies^28,29^ have applied SR101 at a wide range of concentrations (~0.1-500 μM), under which conditions astrocytes stay healthy, and we found 100 nM optimal for our SM*d*M experiments. SM*d*M of the labeled cells was carried out in Leibovitz’s L-15 medium (21083027, Gibco) with 20 mM HEPES (15630106, Gibco).

### Graphene electroporation-based intracellular delivery and SM*d*M in A549 cells

Graphene electroporation was performed as described previously.^30^ Briefly, monolayer graphene (Graphene Supermarket) was transferred onto #1.5 glass coverslips. A549 cells (University of California Berkeley Cell Culture Facility) were cultured following standard protocols in Dulbecco’s Modified Eagle’s Medium (DMEM) supplemented with GlutaMAX-I (10566-016, Gibco) and 10% fetal bovine serum, and plated onto the graphene-coated coverslip. After ~24 h, the culture medium was changed to an electroporation buffer (1652676, Bio-Rad) containing the fluorescent probe to be delivered at ~100 nM. Electroporation was achieved by applying a 15 V voltage pulse (~5-10 ms halftime) across the graphene-coated substrate and a counter electrode ~500 μm above. After ~10 min incubation, the sample was rinsed with the imaging buffer (L-15 medium + 20 mM HEPES) 3 times, and SM*d*M was performed as described above.

### Data analysis

SM*d*M data were analyzed as described previously.^31^ Briefly, single molecules were super-localized in each recorded frame, and their two-dimensional displacements between paired frames were calculated. The accumulated displacement values were spatially binned onto a 320 × 320 nm^2^ grid. For each spatial bin, the distribution of local displacements was fitted through maximum likelihood estimation (MLE) to a probability model based on two-dimensional random walk:^31^

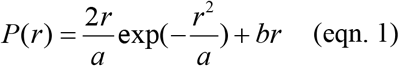

where *r* is the single-molecule displacement in the fixed time interval Δ*t*, *a* = 4*D*Δ*t*, and *b* is a background term accounting for mismatched molecules that randomly diffuse into the field of view during Δ*t*, as discussed previously.^31^ Performing MLE for all spatial bins generated a map of local *D* for color rendering. Analysis of twodimensional vectorial single-molecule displacements showed no noticeable anisotropy, and so fitting to a onedimensional random-walk model^10^ yielded comparable *D* values (**Figure S4**).

## Supporting Information

Oscilloscope-measured timing waveforms; SM*d*M results of Cy3B diffusing in PBS containing different amounts of glycerol; SM*d*M for a control astrocyte not loaded with SR101 but expressing the mEos3.2 FP; Two-dimensional vectorial single-molecule displacement analysis (PDF)

## Acknowledgments

We acknowledge support by the National Institute of General Medical Sciences of the National Institutes of Health (DP2GM132681), the Packard Fellowships for Science and Engineering, and the Heising-Simons Faculty Fellows award.

