## Supplementary Figures for "Single-molecule displacement mapping indicates unhindered intracellular diffusion of small (<~1 kDa) solutes"

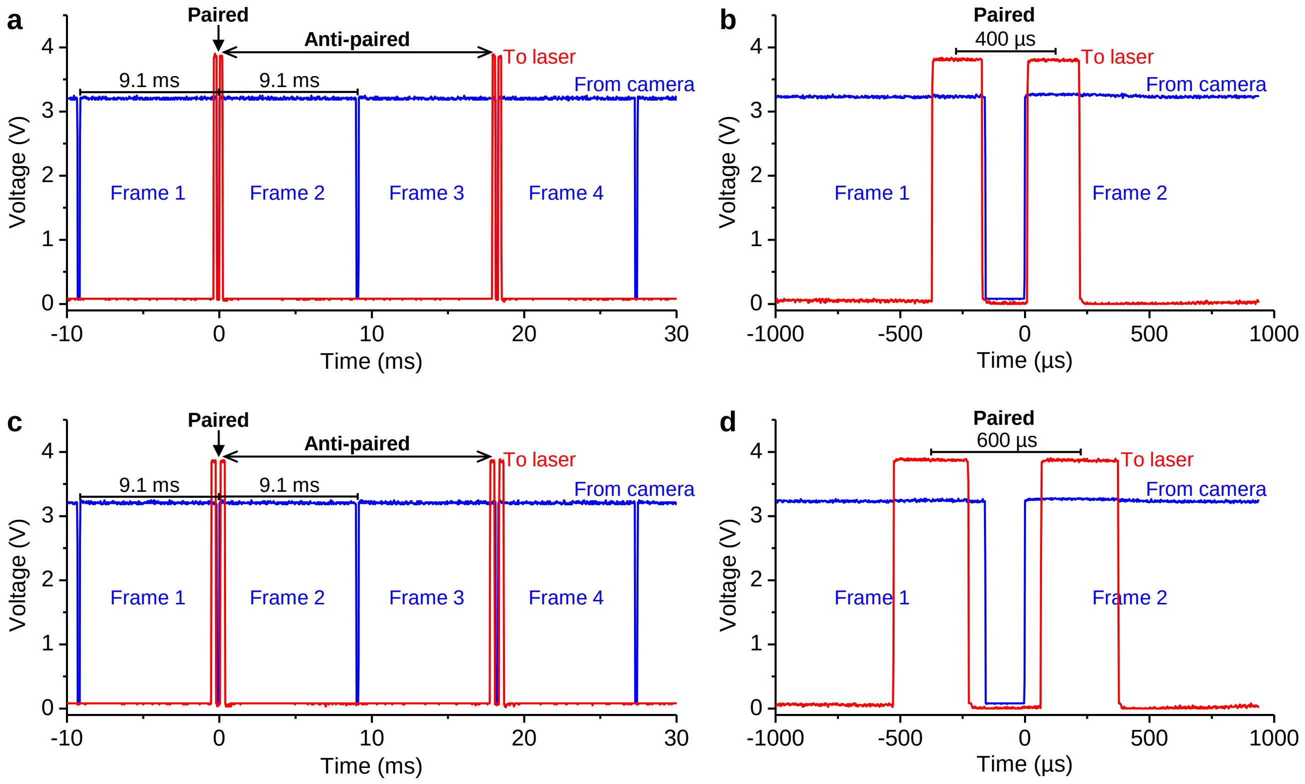


**Figure S1.** Oscilloscope-measured typical timing waveforms in this study. Blue traces: the "fire" TTL signal from the EM-CCD camera, which ran at 110 fps (9.1 ms/frame). 3.3 V: exposure; 0 V: dead time between frames (~160 µs). Red traces: output from the PCI-6733 card, which modulates the 561- or 642-nm excitation lasers between off (low voltage) and on (high voltage) states. (a,b) Pulse center-to-center separation $\Delta t$ = 400 µs and pulse durations $\tau$ = 200 µs, shown as zoom-out for four consecutive camera frames (a) and zoom-in for the tandem pulses across the two paired frames (b). (c,d) Pulse center-to-center separation $\Delta t$ = 600 µs and pulse durations $\tau$ = 300 µs. Note the very long (~18 ms) time separations between the anti-paired pulses.

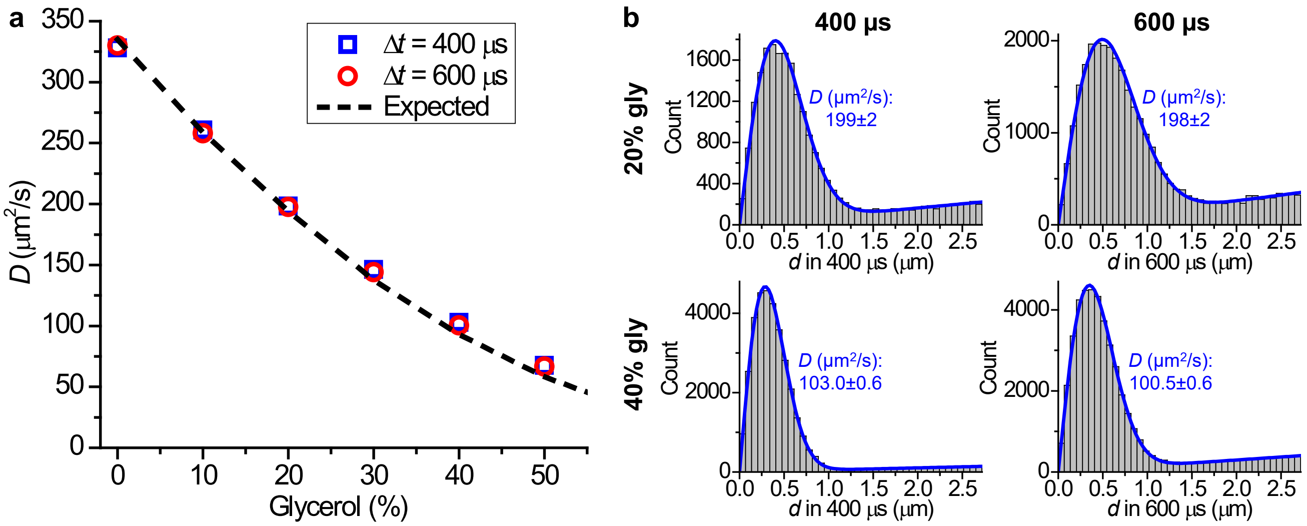


**Figure S2.** SM*d*M results on Cy3B diffusing in PBS containing different amounts of glycerol. (a) SM*d*M-determined diffusion coefficient *D* of Cy3B as a function of the glycerol weight percentage, for experiments carried out at center-to-center pulse separations $\Delta t$ of 400 μs (blue squared) and 600 μs (red circles), compared to that is predicted by scaling the 0% glycerol value with the known glycerol concentration-dependent viscosity (dash line).[1] (b) Distributions of the recorded single-molecule displacements for results in 20% and 40% glycerol at $\Delta t$ = 400 and 600 μs. Blue lines: maximum likelihood estimation (MLE) fits to our model, with resultant *D* values and uncertainties marked in each plot.

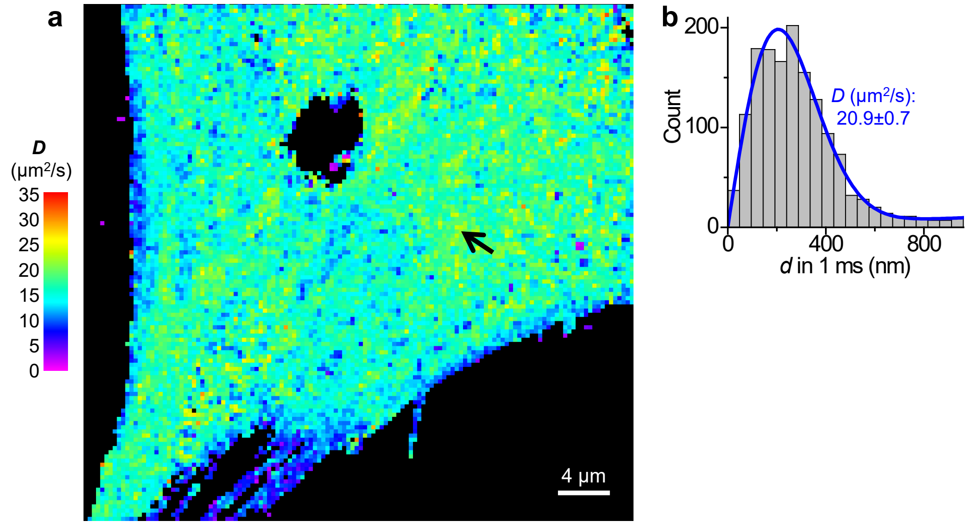


**Figure S3.** SM*d*M under the same 561-nm excitation laser for an astrocyte not loaded with SR101 but expressing the mEos3.2 FP. (a) Color-coded SM*d*M *D* map. Note the 10× lower *D* values for the color scale compared to Figure 2d. (b) Local distribution of the recorded 1-ms single-molecule displacements for an ~1 $\mu m^2$ region pointed to by the black arrow in (a). Blue line: MLE fit to our model, yielding $D = 20.9 \pm 0.7\ \mu m^2/s$.

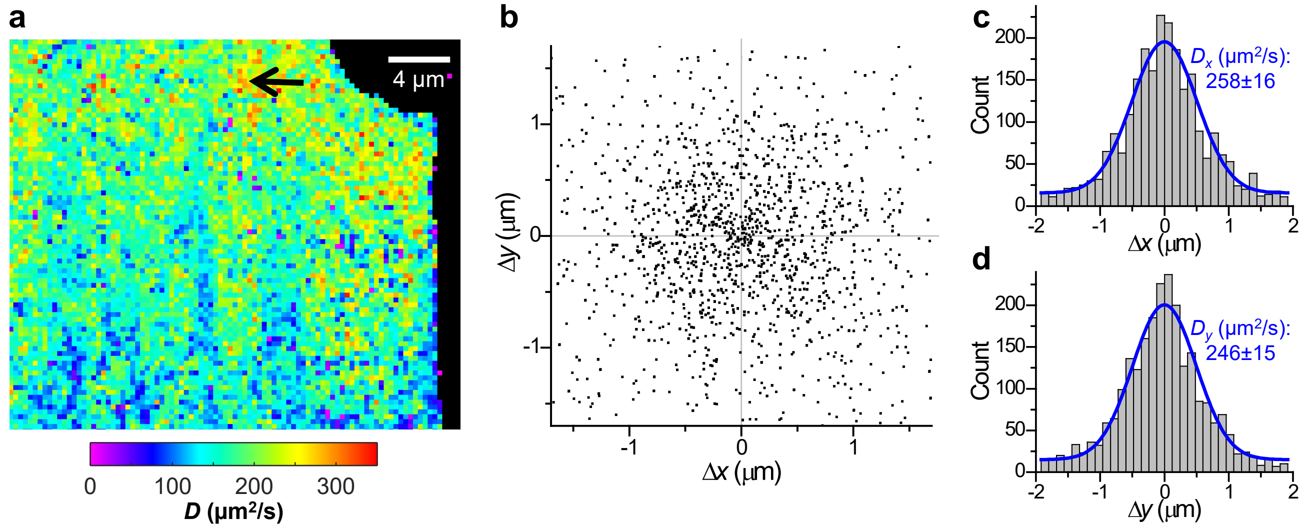


**Figure S4.** Two-dimensional vectorial single-molecule displacement analysis. (a) Color-coded principal-direction SM*d*M (pSM*d*M) diffusivity map for the same data shown in Figure 2d, obtained by first calculating the preferred diffusion direction in each spatial bin, and then fitting the distribution of single-molecule displacements along that direction to a modified one-dimensional random-walk model with the probability distribution of $P(x) = \frac{1}{\sqrt{a\pi}}\exp(-\frac{x^2}{a}) + b$, where $a = 4D\Delta t$ ($\Delta t$ = 500 μs) and $b$ accounts for a uniform background.[2] (b) Two-dimensional plots of the vectorial 500-μs single-molecule displacements accumulated for a region pointed by the black arrow in (a), showing no noticeable anisotropy. (c,d) Distributions of the displacements in (b) along the horizontal (c) and vertical (d) directions. Blue curves: MLE fits to the modified one-dimensional random-walk model above, yielding $D_x$ = 258±16 and $D_y$ = 246±15 μm²/s, comparable to $D$ obtained from fitting the scalar magnitude of single-molecule displacements to the two-dimensional random-walk model (Figure 2e).
